# Ccr4-Not complex reduces transcription efficiency in heterochromatin

**DOI:** 10.1101/2021.08.08.455218

**Authors:** Pablo Monteagudo, Cornelia Brönner, Parastou Kohvaei, Haris Amedi, Stefan Canzar, Mario Halic

## Abstract

Heterochromatic silencing is thought to occur through a combination of transcriptional silencing and RNA degradation, but the relative contribution of each pathway is not known. In this study we analyzed RNA Polymerase II (RNA Pol II) occupancy and levels of nascent and steady-state RNA in different strains of fission yeast, in order to quantify the contribution of each pathway to heterochromatic silencing. We found that transcriptional silencing consists of two components, reduced RNA Pol II accessibility and, unexpectedly, reduced transcriptional efficiency. Heterochromatic loci showed lower transcriptional output compared to euchromatic loci, despite the presence of comparable amounts of RNA Pol II in both types of regions. We determined that the Ccr4-Not complex and H3K9 methylation are required for reduced transcriptional efficiency in heterochromatin and that a subset of heterochromatic RNA is degraded more rapidly than euchromatic RNA. Finally, we quantified the contribution of different chromatin modifiers, RNAi and RNA degradation to each silencing pathway. Our data show that several pathways contribute to heterochromatic silencing in a locus-specific manner and reveal transcriptional efficiency as a new mechanism of silencing.

## Introduction

Heterochromatin is essential to maintain genome stability and transcriptional regulation. Defects in heterochromatin formation lead to aberrant centromere and telomere function, aneuploidy and cancer. In fission yeast, constitutive heterochromatin is established at centromeres, subtelomeres and the silent mating type (*mat*) locus (Allshire and Ekwall, 2015). At centromeric repeats, RNAi is essential for heterochromatin formation as Argonaute, guided by small RNAs, recruits the H3K9 methyltransferase complex CLRC to chromatin (Halic and Moazed, 2010; Holoch and Moazed, 2015; Marasovic et al., 2013; Martienssen and Moazed, 2015; Ugolini and Halic, 2018; Verdel et al., 2004; Volpe et al., 2002). This leads to deposition of repressive H3K9 methylation (H3K9me) mark by Clr4, recruitment of HP1 proteins Chp2 and Swi6, and heterochromatin formation (Allshire and Ekwall, 2015; Holoch and Moazed, 2015; Martienssen and Moazed, 2015).

Current data suggest that heterochromatic silencing occurs through a combination of transcriptional silencing and RNA degradation. At the level of transcriptional silencing, HP1 proteins bind H3K9me nucleosomes and recruit downstream-acting complexes (Castel and Martienssen, 2013; Motamedi et al., 2008; Zocco et al., 2016). HP1 protein Chp2 recruits the complex SHREC to deacetylate chromatin and to remodel nucleosomes in heterochromatin, which is required for silencing (Creamer et al., 2014; Motamedi et al., 2008; Sugiyama et al., 2007). These activities were suggested to reduce RNA Pol II access (Chen and Widom, 2005; Schuettengruber et al., 2007).

Heterochromatin is also thought to promote recruitment of the RNA degradation machinery to degrade nascent transcripts (Brönner et al., 2017; Bühler et al., 2008; Cotobal et al., 2015; Marasovic et al., 2013; Pisacane and Halic, 2017; Reyes-Turcu et al., 2011; Reyes-Turcu and Grewal, 2012; Sugiyama et al., 2016). In fission yeast, RNA Pol II-transcribed heterochromatic transcripts are polyadenylated (pA) products and undergo degradation by the RNAi pathway, by the Ccr4-Not complex and by the exosome (Brönner et al., 2017). The first step of mRNA degradation is generally shortening of the 3’ pA tail by the Ccr4-Not complex and Pan nucleases (Wahle and Winkler, 2013). This induces removal of the 5’ cap which enables 5’-3’ degradation by the exonuclease Xrn1 (Exo2 in *S. pombe*) and 3’-5’ degradation by the exosome subunit Rrp6 (Houseley et al., 2006).

How transcriptional silencing and RNA degradation pathways collaborate and their relative con contributions to silencing is not known. In this study, we quantified the contribution of these pathways to heterochromatic silencing by analyzing RNA Pol II occupancy, nascent RNA and steady-state RNA in different fission yeast strains. We also defined the heterochromatic factors that contribute to transcriptional silencing and/or RNA degradation. We found that transcriptional silencing occurs through reduced RNA Pol II accessibility, as previously proposed, but, unexpectedly, also through reduced transcriptional efficiency, a mechanism that had not been previously described. Our data revealed that RNA Pol II transcriptional output is lower at heterochromatic loci compared to euchromatin loci with the same levels of RNA Pol II occupancy. We determined that the Ccr4-Not complex and H3K9 methylation are essential for the reduced transcriptional efficiency at heterochromatin and quantified the contributions of heterochromatin factors to the reductions in RNA Pol II occupancy, transcriptional efficiency and RNA stability.

## Results

### Transcriptional silencing and degradation of heterochromatic RNA

To dissect the contributions of transcriptional silencing and RNA degradation to heterochromatic silencing, we analyzed RNA Pol II occupancy, nascent RNA, and steady-state RNA at heterochromatic and euchromatic regions in fission yeast (**Figure 1A** shows data for specific genomic regions from wild-type cells). For Pol II occupancy, we performed ChIP-seq analyses of serine 2 phosphorylated (S2P)-RNA Pol II-bound DNA. To reduce the noise in our data analysis, we subtracted normalized input from all RNA Pol II ChIP-seq datasets. For the normalization, the background is defined as the centromeric central core (Cenp-A containing chromatin), a region with the lowest number of reads in RNA Pol II ChIP-seq; we thus subtracted input so that this region would show near zero reads (**Supplemental Figure S1A, see also Methods**). For nascent RNA, we performed RNA-seq of S2P-RNA Pol II-bound RNA (Pol II RIP). Quality of nascent RNA was assessed by retention of intronic sequences, which are strongly enriched in nascent RNA (**Supplemental Figure S1B, C**). For steady-state RNA we sequenced polyadenylated (pA) RNA and total RNA. Both datasets showed comparable levels of heterochromatic transcripts, indicating that these are mostly polyadenylated (**Supplemental Figure S1D and E**).

**Figure 1.**
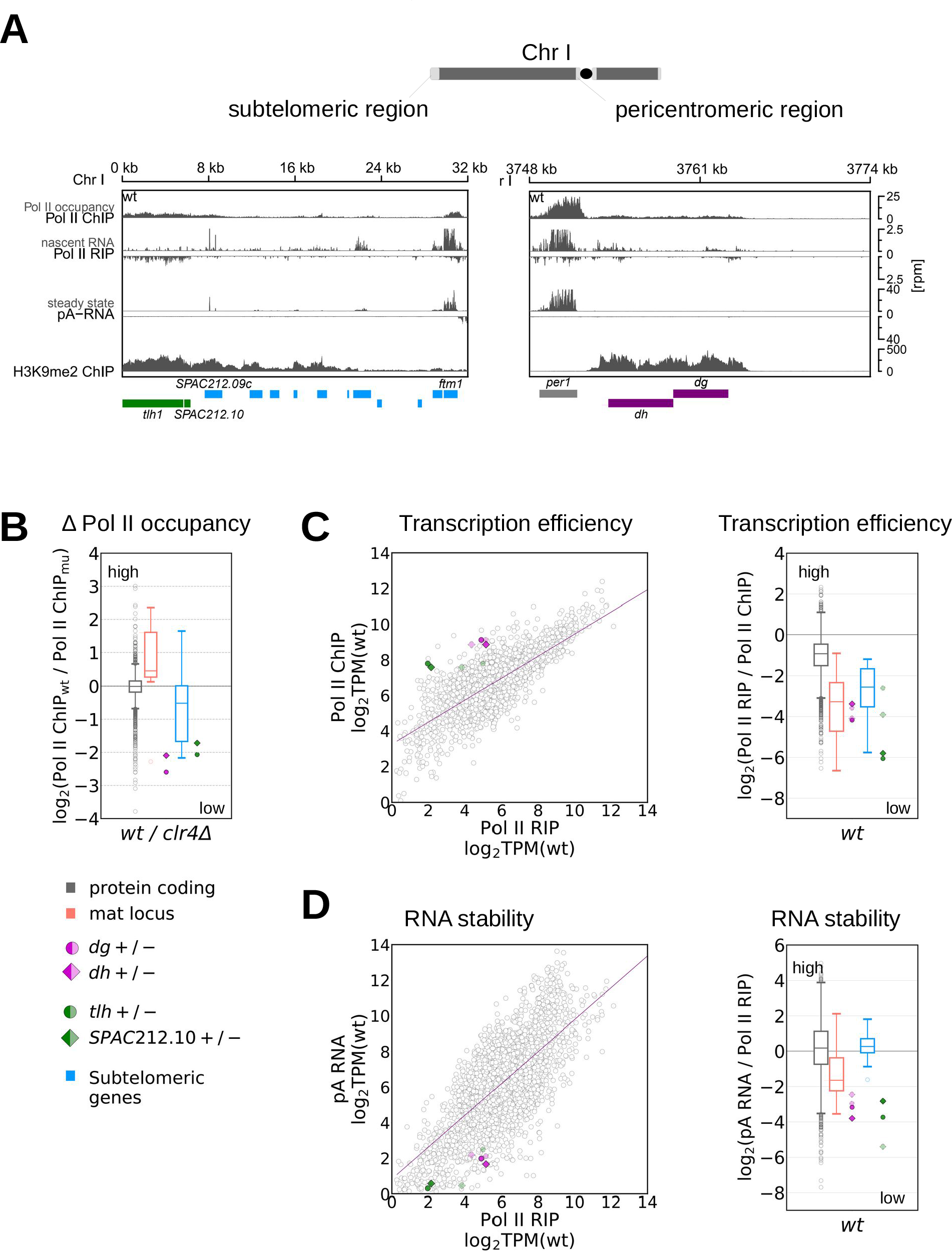
Heterochromatic repeats have reduced RNA Pol II occupancy, Transcription Efficiency and RNA stability. (**A**) Analysis of the next-generation sequencing data showing occupancy of S2-phosphorylated RNA Pol II (ChIP-seq), nascent RNA (S2P-Pol II RIP-seq), steady state RNA levels (pA RNA-seq) and H3K9me2 levels (ChIP-seq) at subtelomeric and pericentromeric regions in *S. pombe* wild-type cells. Gene locations are indicated as boxes below the coverage and color-coded: gray, protein-coding genes; purple, centromeric *dg, dh*; green, subtelomeric loci *tlh* and *SPAC212.10*; blue, other subtelomeric genes. (**B**) Box plot showing RNA Pol II occupancy (S2P-Pol II ChIP-seq) in wild-type cells relative to *clr4Δ* cells for protein-coding genes (gray), mat locus (orange), and subtelomeric genes (blue). For protein-coding genes, individual transcript are shown as circles, bottom and top of the box correspond to lower and upper quartiles of the data, bar is the median and whiskers are median +/-1.5 times interquartile range. Colored symbols on the right show centromeric *dg* and *dh* (dark purple), subtelomeric *tlh and SPAC212.10* (dark green). Each data point is the average of at least two independent samples. (**C**) Transcription efficiency in wild-type cells. Left, S2P-Pol II ChIP-seq (Pol II occupancy) data plotted over S2P-Pol II RIP-seq data (nascent RNA). TPM, transcripts per million. Gray circles are individual protein-coding genes; regression line is also shown in gray. Also plotted are centromeric *dg* and *dh* (dark purple for + strand, bright purple for -strand) and *tlh* and *SPAC212.10* (dark green for + strand, bright green for - strand). Each data point is the average of at least two independent samples. Right, box plot showing transcription efficiency distributions by gene categories, data are plotted and color-coded as in panel B. (**D**) RNA stability in wild-type cells. Left, pA RNA-seq (steady-state RNA) data plotted over S2P-Pol II RIP seq data (nascent RNA). Data are plotted as defined for panel C. Right, box plot showing RNA stability (pA RNA / Pol II RIP) distribution by gene categories, data are plotted and color-coded as in panel B.

To determine how heterochromatin changes RNA Pol II accessibility, we compared RNA Pol II occupancy in wild-type cells with the occupancy in cells having a deletion of the H3K9 methyltransferase Clr4, and thus do not have H3K9me and heterochromatin (**Figure 1A, B**). We did not detect substantial changes in RNA Pol II occupancy at protein-coding genes between wild-type and clr4Δ cells. At heterochromatic regions, we observed different effects according to the specific locus. At centromeric *dg*/*dh* and subtelomeric *tlh* repeats (*tlh* and SPAC212.10), RNA Pol II occupancy was ∼four-fold lower in wild-type cells compared to *clr4Δ* cells (**Figure 1A, B**). At other subtelomeric regions or at the mat locus, we did not observe a substantial change in RNA Pol II occupancy in the absence of heterochromatin. (**Figure 1B**).

Our data also showed that RNA Pol II complexes can be present in heterochromatic regions, but they do not always actively transcribe or produce nascent RNA. For example, in wild-type cells, heterochromatic *tlh1* and subtelomeric *ftm1* loci show similar RNA Pol II occupancy, but *ftm1* produces substantially more nascent RNA (**Figure 1A**). To analyze this relationship further, we plotted the nascent RNA levels over chromatin-bound RNA Pol II for individual loci in wild-type cells; we also calculated the ration between those measurements, which informs on how much nascent RNA is synthesized by RNA Pol II at any given locus. We define this parameter as transcription efficiency (**Figure 1C**).

We found a linear correlation between RNA Pol II occupancy and nascent RNA when examining ∼5 000 euchromatic, protein-coding loci in wild-type cells, thus transcription efficiency was constant across the examined euchromatic loci. In contrast, heterochromatic loci showed lower transcriptional efficiency (8-fold on average) compared to protein-coding (**Figure 1C**). This observation applied to all heterochromatic regions in fission yeast. Thus, in wild-type cells, heterochromatic loci are less efficiently transcribed than protein-coding loci that have the same amount of RNA Pol II, indicating that RNA Pol II does not productively transcribe heterochromatic regions.

Next, we calculated the ratio of steady-state RNA to nascent RNA, which informs on the stability of that specific transcript. Notably, we found that a subset of heterochromatic transcripts are less stable than the average transcript from a protein-coding locus (**Figure 1E and F**). This subset of unstable heterochromatic RNAs includes transcripts from centromeric *dg/dh* and subtelomeric *tlh* regions, which had been previously shown to be degraded by RNAi and by the Ccr4-Not complex, respectively (Brönner et al., 2017; Halic and Moazed, 2010; Marasovic et al., 2013). For transcripts derived from the other subtelomeric genes or from the *mat* locus, RNA stability was comparable to transcripts from protein-coding genes (**Figure 1F**), indicating that those loci primarily undergo transcriptional silencing, with no major role for RNA degradation in their silencing.

### Effect of heterochromatin on silencing pathways

Our data indicate that tree different pathways can contribute to heterochromatin silencing: RNA Pol II occupancy, transcriptional efficiency and RNA degradation. To further examine the impact of heterochromatin structure on those pathways, we compared the contributions from each of them at heterochromatic loci from wild-type and *clr4Δ* cells.

We found that RNA Pol II transcribes repetitive regions more efficiently in the *clr4Δ* cells compared to wild-type cells. RNA Pol II occupancy at centromeric *dg/dh* and subtelomeric *tlh* repeats was ∼four-fold higher in *clr4Δ* cells relative to wild-type cells, the increase in nascent RNA was ∼twenty-fold (**Figure 2A and B**). The data also demonstrate that heterochromatin structure contributes to reduced transcriptional efficiency: in *clr4Δ* cells, all heterochromatic regions (centromeres, subtelomeres and the mat locus) reach transcriptional efficiency that is comparable to euchromatic regions (**Figure 2C**). Moreover, transcription efficiency is the single mode of heterochromatic silencing that acts in all heterochromatic regions.

**Figure 2:**
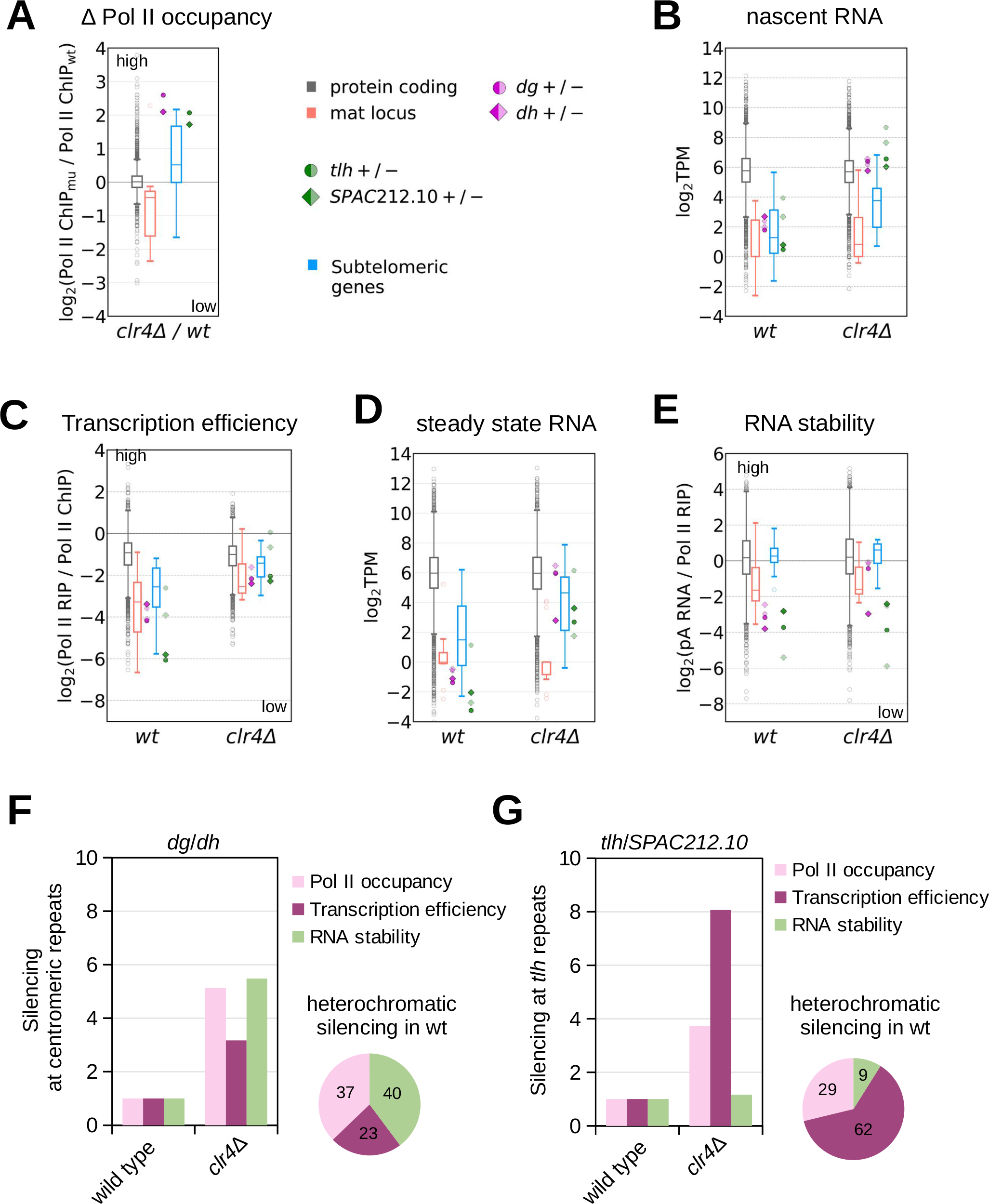
Contribution of distinct pathways to heterochromatic silencing. (**A**) RNA Pol II occupancy (Pol II ChIP-seq) in clr4Δ cells relative to wild-type cells for indicated gene categories; data are the same as in Figure 1B, with inverted ratios. (**B**) Nascent RNA (Pol II RIP) in wild type and *clr4Δ* cells. Data are plotted as defined for Figure 1B. (**C**) Transcription efficiency (Pol II RIP / Pol II ChIP) in wild-type and *clr4Δ* cells. Data are plotted as defined for Figure 1C. (**D**) Steady state RNA (pA RNA-seq) shown for wild-type and *clr4Δ* cells. Data are plotted as defined for Figure 1B. (**E**) RNA stability (pA RNA / Pol II RIP) in wild type and *clr4Δ* cells. Data are plotted as defined for Figure 1D. (**F, G**) Bar graphs showing loss of silencing at the three pathways (Pol II occupancy, transcription efficiency and RNA stability) at centromeric dg and dh (D) and at subtelomeric tlh (E) in clr4Δ cells, relative to wild type. Pie charts show contribution of each pathways to heterochromatic silencing at repeats in wild-type cells. Average of at least two independent samples is shown for all figures.

We next assessed RNA stability in the absence of heterochromatin structure. We observed that subtelomeric tlh RNA has similarly low stability in *clr4Δ* compared to wild-type cells, indicating that multiple pathways degrade RNA from subtelomeric *tlh* repeats, independently of heterochromatin structure, in agreement with previous observations (Brönner et al., 2017). In contrast, centromeric *dg/dh* transcripts showed increased stability in *clr4Δ* compared to wild-type cells, indicating that heterochromatin structure is required to target those transcripts to degradation (**Figure 2D and E**).

These data with clr4Δ confirm our observations with wild-type cells: at centromeric dg/dh, silencing results from a combination of reduced RNA Pol II occupancy (Figure 2A), reduced transcriptional efficiency (**Figure 2C**) and increased RNA degradation (**Figure 2E**), whereas for subtelomeric tlh repeats, silencing occurs primarily at the transcriptional level.

By comparing data from wild-type and *clr4Δ* cells, we quantified the relative contribution of each pathway to heterochromatic silencing. In wild-type cells, RNA degradation and transcriptional silencing contribute similarly to silencing of centromeric repeats (40% and 60%, respectively) (**Figure 2F**) with transcriptional silencing further parsed out into RNA Pol II occupancy (37%) and transcriptional efficiency (23%). At subtelomeric *tlh* repeats, heterochromatic silencing occurs primarily by transcriptional silencing (92%) (**Figure 2G**), which is predominately mediated through reduced transcriptional efficiency (62%). A similar pattern is observed at the remaining subtelomeric regions and the mat locus: silencing is primarily transcriptional (∼80%), with transcriptional efficiency (∼50%) being the major silencing pathway (**Supplemental Figure S1F, G**).

### Contribution of heterochromatin factors to distinct silencing pathways

Next, we analyzed RNA Pol II occupancy, transcriptional efficiency and RNA degradation in strains defective in the RNAi machinery (*ago1Δ*), lacking chromatin modifiers (*clr3Δ*, *mit1Δ*, *chp2Δ* and swi6Δ) or RNA degradation components (*rrp6Δ*, *exo2Δ, caf1Δ, ccr4Δ* and *mot2Δ*) (**Supplemental Figure S3-S6**). Chp2 and Swi6 are HP1 family proteins; wherein Chp2 recruits the SHREC complex to chromatin. SHREC subunits, Mit1 and Clr3, are a chromatin remodeler and a histone deacetylase, respectively (Motamedi et al., 2008; Sugiyama et al., 2007). Rrp6 is a component of the exosome complex; Exo2 is a 5’->3’ exonuclease. Caf1, Ccr4 and Mot2 are components of Ccr4-Not deadenylase complex.

Our data show that reduced RNA Pol II occupancy at centromeric repeats requires RNAi (**Figure 3A** **and Supplemental Figure S2A, S3A-C**), several chromatin modifiers (**Figure 3A** **and Supplemental Figure S3D-E, S4A-B**) and the RNA degradation machinery (**Figure 3A** **and Supplemental Figure S4C-D**). At subtelomeric *tlh* repeats, only chromatin modifiers are required to limit RNA Pol II occupancy (**Figure 3A** **and Supplemental Figure S2B, S3D-E, S4A-B**). At the remaining subtelomeric genes and at the *mat* locus, we did not observe strong increase in RNA Pol II occupancy in absence of heterochromatin factors (**Figure 3A, B** **and Supplemental Figure S2C, D**). Finally, the Ccr4-Not complex components showed little effect on RNA Pol II occupancy at all heterochromatic regions (**Figure 3B** **and Supplemental Figure S5A-C**).

**Figure 3:**
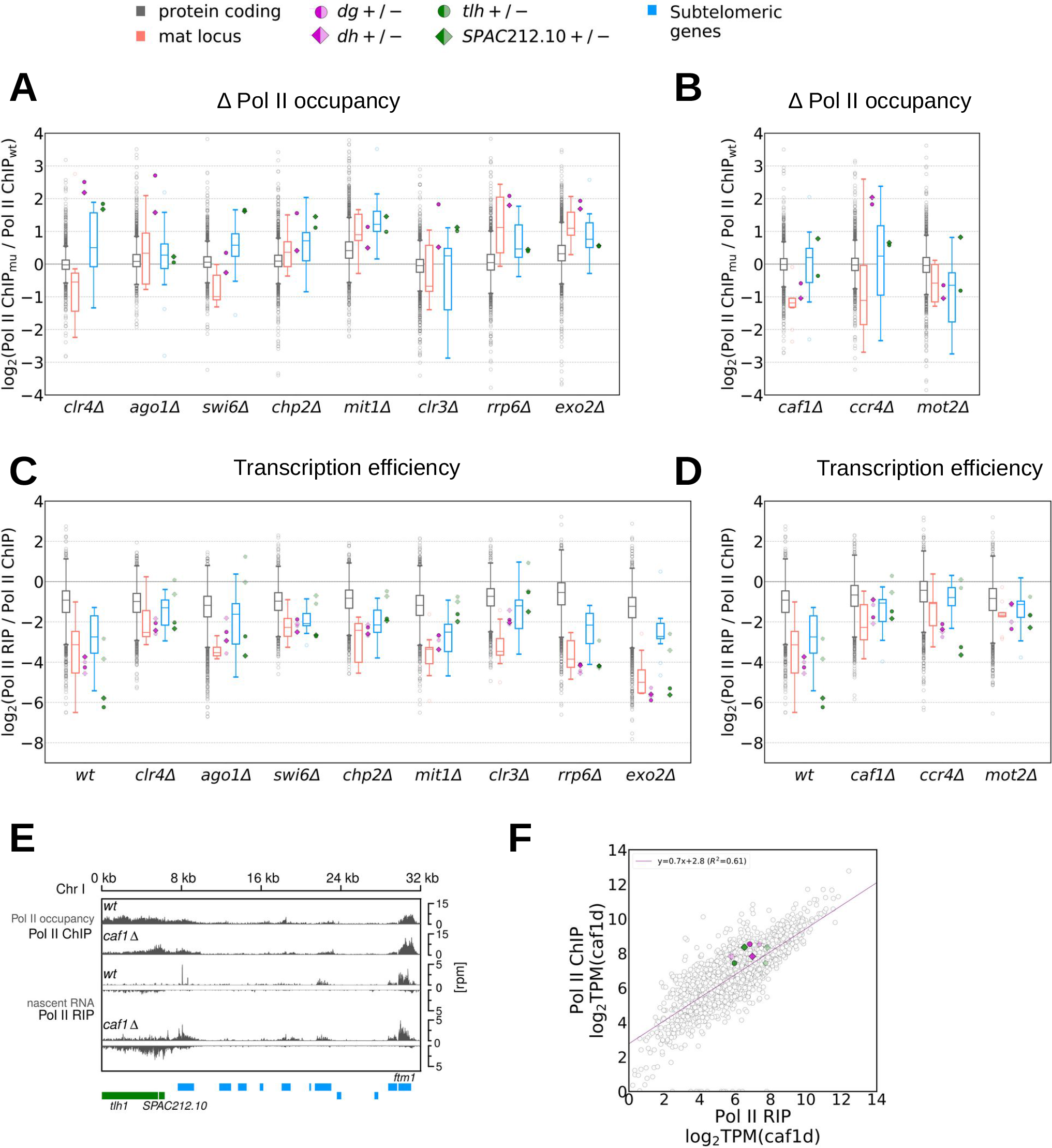
RNA Pol II occupancy, transcription efficiency and RNA stability in different mutants. (**A, B**) Box plots showing ration of RNA Pol II occupancy (S2P-Pol II ChIP-seq) in mutants compared to wild type over indicated genes. Shown are mutants in factors involved in heterochromatin formation and RNA degradation (A) or in Ccr4-Not complex components (B). Data are plotted as defined for Figure 1B. (**C, D**) Box plot showing transcription efficiency (Pol II RIP / Pol II ChIP) over indicated genes in wild type and mutants in factors involved in heterochromatin formation and RNA degradation (C) or in Ccr4-Not complex components (D). Data are plotted as defined for Figure 1B. (**E**) Analysis of the next-generation sequencing data showing occupancy of S2-phosphorylated RNA Pol II (ChIP-seq), nascent RNA (S2P-Pol II RIP-seq), steady state RNA levels (pA RNA-seq) and H3K9me2 levels (ChIP-seq) at subtelomeric and pericentromeric regions in *S. pombe* wild-type and *caf1Δ* cells. Gene locations are indicated as boxes below the coverage and color-coded: gray, protein-coding genes; purple, centromeric *dg, dh*; green, subtelomeric loci *tlh* and *SPAC212.10*; blue, other subtelomeric genes. (**F**) Transcription efficiency in *caf1Δ* cells. S2P-Pol II ChIP-seq (Pol II occupancy) data plotted over S2P-Pol II RIP-seq data (nascent RNA). TPM, transcripts per million. Gray circles are individual protein-coding genes; regression line is also shown in gray. Also plotted are centromeric *dg* and *dh* (dark purple for + strand, bright purple for - strand) and *tlh* and *SPAC212.10* (dark green for + strand, bright green for - strand). Each data point is the average of at least two independent samples.

The reduced transcriptional efficiency at heterochromatic loci depends on most chromatin modifiers (Clr4, HP1 proteins and SHREC complex), but not on RNA degradation factors Rrp6 and Exo2 (**Figure 3C** **and Supplemental Figure S2-S5**). Those chromatin modifiers seem to reduce transcriptional efficiency more than RNA Pol II occupancy (**Supplemental Figure S2**), an effect similar to what we had observed in *clr4Δ* cells. The strongest effect on transcriptional efficiency in heterochromatic regions was observed for components of the Ccr4-Not complex (**Figure 3D** **and Supplemental Figure S2**). The Ccr4-Not deadenylase complex could affect transcriptional efficiency indirectly through changes in RNA levels of other factors involved in heterochromatin formation.

However, analysis of nascent RNA, total RNA and pA RNA data show that RNA levels of heterochromatin formation factors do not change substantially in *caf1Δ* cells compared to wild-type cells (**Supplemental Figure S6E**). This suggests a direct effect of Ccr4-Not on transcriptional efficiency.

Although RNA Pol II occupancy at *tlh* repeats is comparable in *caf1Δ* and wild-type cells, *caf1Δ* cells produce ∼10 fold more nascent RNA from that locus than in wild-type cells (**Figure 3E, F**). This increases transcriptional efficiency in heterochromatic regions in *caf1Δ* cells, which is now comparable to protein-coding genes (**Figure 3D-F**). Notably, H3K9me levels are mostly unaffected at *tlh* and centromeric *dg/dh* repeats in *caf1Δ* cells (**Supplemental Figure S6F**), indicating that increased transcriptional efficiency does not interfere with H3K9me and heterochromatin. These data show that Caf1 reduces transcriptional efficiency after H3K9me is deposited and heterochromatin is established; however, Caf1 requires H3K9me and heterochromatin for its activity, as seen by our data on *clr4Δ* (**Figure 2C**). Thus, the Ccr4-Not complex is required to reduce transcriptional efficiency in heterochromatin regions and it modulates RNA Pol II activity in a heterochromatin-dependent way.

Our data reveal that RNA degradation primarily contributes to silencing of centromeric *dg* and *dh* transcripts (**Figure 4A** **and Supplemental Figure S2A**). This requires RNAi and H3K9me (Clr4), but not other heterochromatic or the RNA degradation factors examined. Of the latter, only deletion of *exo2* showed a small decrease in RNA degradation at the centromeric region, suggesting that multiple RNA degradation pathways act redundantly to degrade heterochromatic transcripts (**Figure 4A** **and Supplemental Figure S2A**). In contrast, deficiency in the Ccr4-Not complex components increased degradation of heterochromatic transcripts compared to wild-type cells; this effect is consistent with increased transcriptional efficiency in this mutant, leading to higher amounts of nascent RNA (**Figure 4B** **and Supplemental Figure S2, S7A**). This increased transcriptional output at heterochromatic regions targets the RNAi machinery to these regions, thus leading to increased degradation (Brönner et al., 2017).

**Figure 4:**
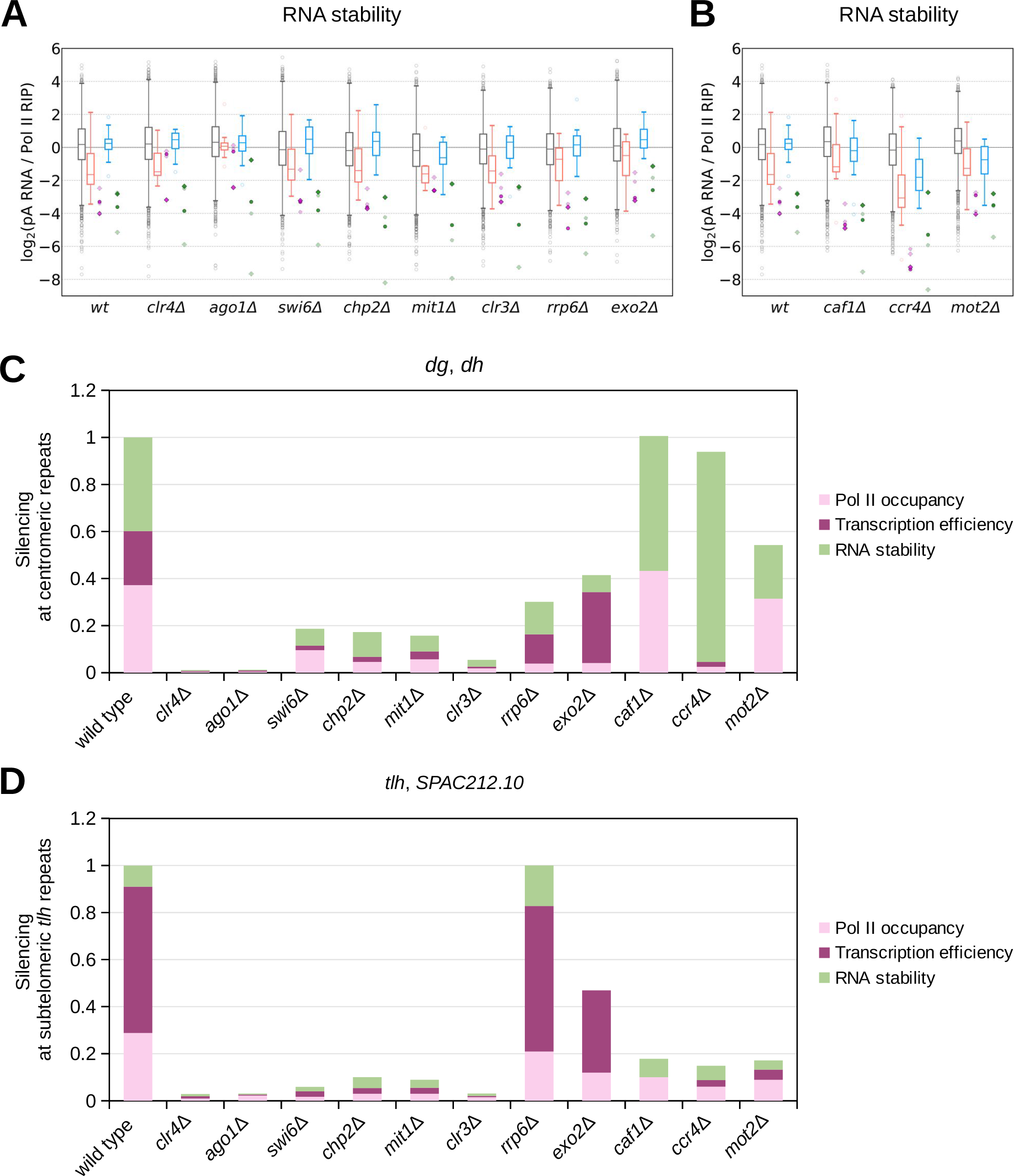
Contribution of individual factors to each pathway of heterochromatic silencing. (**A, B**) Box plot showing RNA stability (pA RNA / Pol II RIP) over indicated genes in wild type and mutants in factors involved in heterochromatin formation and RNA degradation (A) or in Ccr4-Not complex components (B). Data are plotted as defined for Figure 1B. (**C, D**) Bar charts displaying contribution of each of the pathways that are still active in the mutants to silencing at centromeric dg/dh regions (C) and subtelomeric tlh regions (D).

### Contribution of individual proteins to the heterochromatic silencing pathways

For all the mutant strains examined here we calculated the contribution of those factors to each of the three silencing pathways at different heterochromatic loci. Quantification of each pathway’s contribution to heterochromatic silencing was calculated relative to *clr4Δ*, which was defined as a complete loss of heterochromatic silencing. We observed that Ago1 is essential for all three silencing pathways at the centromeric region; in fact, heterochromatin silencing is completely lost in strains lacking those factors (**Figure 4C, D** **and Supplemental Figure S7B, C**). In the absence of the chromatin modifiers silencing overall was strongly reduced, but each pathway was still active and contributed to the remaining silencing (**Figure 4C, D** **and Supplemental Figure S7B, C**). In contrast, deletions of RNA degradation factors led to more limited loss of silencing, with varying effects between centromeric and subtelomeric regions.

We had previously shown that the Ccr4-Not complex degrades subtelomeric RNA redundantly with RNAi (Brönner et al., 2017). Our new data show that the Ccr4-Not complex is also required for silencing at the transcriptional level, where it specifically regulates RNA Pol II transcriptional efficiency at all loci (**Figure 4C, D** **and Supplemental Figure S2**). In agreement with our previous data, deletion of *caf1* had no or little effect on RNA stability and RNA Pol II occupancy, but we show here that it is essential for reduced transcriptional efficiency. Although transcriptional efficiency is increased in *caf1Δ, ccr4Δ* and *mot2Δ* cells, the overall loss of silencing at centromeric repeats is small and increased transcriptional efficiency is compensated by reduced RNA Pol II occupancy and increased RNA degradation (**Figure 3B, 4A, B and Supplemental Figure 2**). The loss of silencing in *caf1Δ, ccr4Δ* and *mot2Δ* cells was more pronounced at subtelomeric *tlh* repeats (compared to centromeric loci), since transcriptional efficiency is the dominant silencing pathway, and increased degradation by RNAi (Brönner et al., 2017) is not sufficient to compensate for its loss (**Figure 4C, D**). Notably, at subtelomeric regions other than *tlh* and at the *mat* locus, the transcriptional efficiency pathway remains active in allmost all mutant strains examined; the exception were mutants for the components of the Ccr4-Not complex, in which transcriptional efficiency is abolished, but other pathways were fully functional (**Supplemental Figure S7B, C**). These data show that Ccr4-Not complex specifically affects the transcriptional efficiency pathway.

To determine which factors affect heterochromatin formation in a similar way, we calculated the Pearson correlation between the effects on RNA Pol II occupancy and transcriptional efficiency across all heterochromatic transcripts. In agreement with our previous calculations, the t-distributed stochastic neighbor embedding t-SNE plot showed that the RNA degradation machineries *exo2* and *rrp6* tend to co-localize together. Chromatin modifier fasctors *clr4*, *clr3*, *mit1*, *swi6* and *chp2* showed similar behavior across all heterochromatic genes (**Figure 5A**), as their deletions increased RNA Pol II occupancy and transcriptional efficiency, and had a smaller effect on RNA degradation, indicating that these proteins mainly act in transcriptional silencing. Components of the Ccr4-Not complex *caf1, ccr4* and *mot2* co-localize as well, indicating their specialized role in transcriptional silencing (**Figure 5A**). The Ccr4-Not is required for transcriptional efficiency at all regions, whereas RNAi is specifically required for all types of silencing at centromeric repeats.

**Figure 5:**
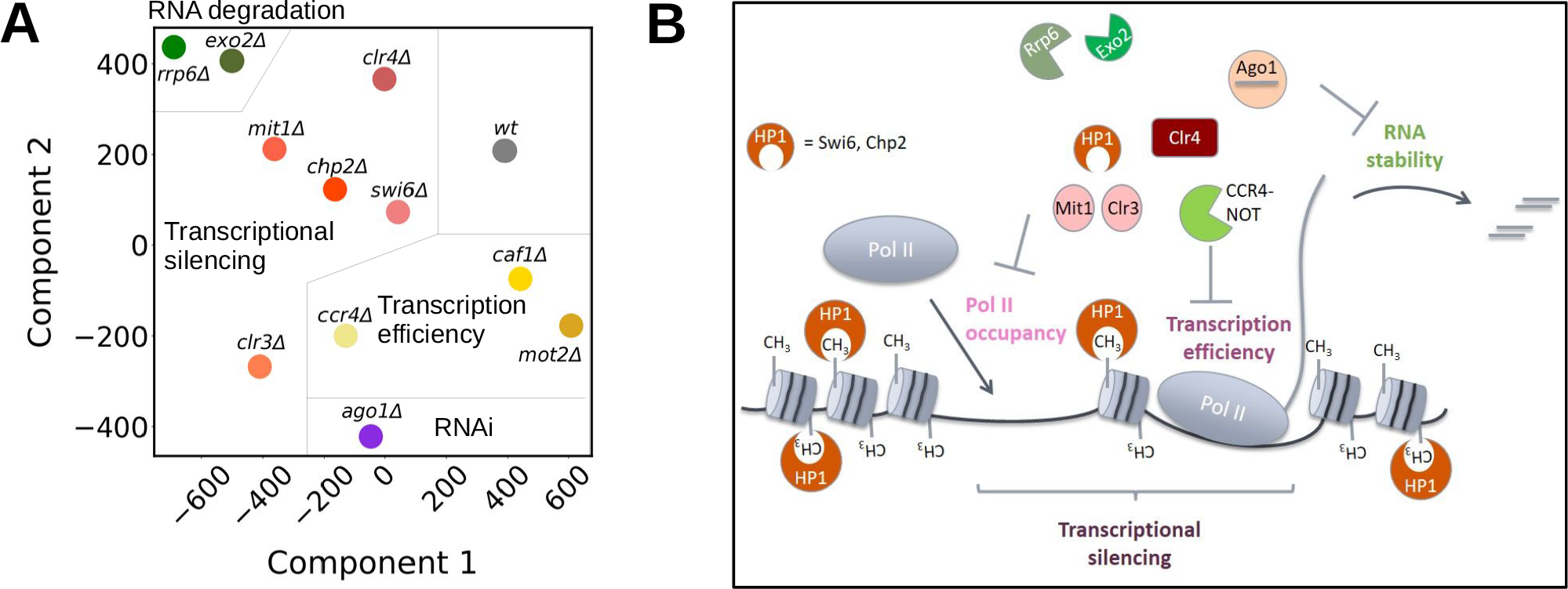
Heterochromatic silencing is a combination of reduced RNA Pol II occupancy, Transcription Efficiency and RNA stability. (**A**) The t-distributed stochastic neighbor embedding (t-SNE) plot showing two dimensional embedding of Pol II occupancy and transcription efficiency. Close proximity of mutants visualizes similarities in transcriptional silencing. (**B**) Schematic presentation of how different proteins involved in heterochromatin formation or RNA degradation contribute to heterochromatic silencing. Three pathways are important for heterochromatic silencing: transcriptional silencing, consisting of Pol II occupancy and transcription efficiency, and RNA degradation.

## Discussion

Our data show that heterochromatic silencing occurs through different mechanisms. First, RNA Pol II accessibility is reduced at heterochromatic regions, which also reduces overall transcription at those loci (**Figure 5B**). Second, we identified transcriptional efficiency as a new mode of heterochromatic silencing; although RNA Pol II is present at heterochromatic loci, the transcriptional efficiency is reduced by heterochromatin and the Ccr4-Not complex (**Figure 5B**). This mode of silencing is present at all heterochromatic regions in fission yeast cells. Third, the final compoment of heterochromatic silencing is RNA degradation by RNAi and several RNA degradation machineries, including the Ccr4-Not complex.

Reduced RNA Pol II access was initially proposed as the major mode of heterochromatic silencing and has been observed in many organisms (Feng and Michaels, 2015; Grewal and Elgin, 2007). In fact, we observed that RNA Pol II occupancy at centromeres strongly increased in cells with mutations that greatly affect H3K9me levels, such as *clr4Δ* or *ago1Δ* (Okita et al., 2019). Components of the heterochromatin pathway that act downstream of H3K9me, such as HP1 proteins, are required to reduce RNA Pol II occupancy, but to a lesser extent than Clr4. Notably, RNA degradation components Exo2 and Rrp6 are required for reduced RNA Pol II occupancy at the centromeric region, in agreement with our previous finding that RNA degradation is required for formation of heterochromatic domains (Brönner et al., 2017).

In addition to reduced RNA Pol II occupancy, we identify reduced transcriptional efficiency as another mode of transcriptional silencing. Although RNA Pol II is present at heterochromatic regions, its RNA production (nascent RNA levels) is proportionally lower than in euchromatic regions. This silencing pathway occurs at all heterochromatic regions in fission yeast cells and might be conserved in other organisms as well. In fact, this mode of silencing might be analogous to SIR-mediated silencing in *S. cerevisiae,* wherein transcription is initiated but elongation is blocked by the SIR complex, which maintains RNA Pol II in a stalled conformation (Johnson et al., 2013).

We found that heterochromatin is essential to reduce transcriptional efficiency: in the absence of H3K9me, transcription efficiency at heterochromatic loci is increased to the level of euchromatic genes. These results also show that reduced transcriptional efficiency is not encoded in the DNA sequence itself, but it is controled by the heterochromatin. Notably, we show that reduced transcriptional efficiency is also mediated by the Ccr4-Not complex, as RNA Pol II occupancy was not altered at heterochromatic regions of *caf1Δ* cells, but transcriptional efficiency increased to the level of protein-coding genes. Notably, H3K9me levels at subtelomeric *tlh* and centromeric *dg*/*dh* repeats were only modestly affected in *caf1Δ* cells (**Supplemental Figure S6F**) (Brönner et al., 2017; Cotobal et al., 2015; Sugiyama et al., 2016), indicating that heterochromatin formation is functional in those cells. Thus, the Ccr4-Not complex regulates transcriptional efficiency post-heterochromatin formation, but H3K9me and heterochromatin are required for this regulation by the Ccr4-Not complex.

Ccr4-Not was initially described as a chromatin-associated complex involved in transcription (Miller and Reese, 2012), and later shown to act as a transcription elongation factor that would reactivate arrested RNA Pol II (Dutta et al., 2015; Kruk et al., 2011). It is plausible, that, in presence of heterochromatic marks H3K9me and histone deacetylation, Ccr4-Not exhibits the opposite effect and stalls RNA Pol II. Recently, Ccr4-Not was shown to be required for DNA-damage dependent ubiquitination and degradation of RNA Pol II (Jiang et al., 2019), and Ccr4-Not mediated stalling could require RNA Pol II ubiquitination. Furthermore, the Ccr4-Not complex is recruited to RNA Pol II by the histone chaperone Spt6 (Dronamraju et al., 2018), which was also implicated in heterochromatin formation in fission yeast (Kiely et al., 2011). Altogether, these various observations support that Ccr4-Not is recruited to chromatin and regulates RNA Pol II transcription.

Our data show that RNA degradation contributes to heterochromatic silencing at centromeric repeats and *tlh*, but not at other subtelomeric genes or at the *mat* locus. RNA degradation at centromeric region is dependent on RNAi and heterochromatin, but those are not required for RNA degradation at subtelomeric *tlh* repeats. This observation is in agreement with the data that, at subtelomeric *tlh* repeats, RNA is degraded by parallel mechanisms that are heterochromatin dependent and independent (Brönner et al., 2017).

In conclusion, we identified a new mode of heterochromatic silencing termed transcriptional efficiency. This mode of silencing depends on H3K9me and the Ccr4-Not complex and acts as a dominant silencing pathway at most heterochromatic loci in fission yeast.

## Data Access

The sequencing data reported in this paper have been submitted to the NCBI Gene Expression Omnibus (GEO; http://www.ncbi.nlm.nih.gov/geo/) under the accession number **XXXXX**.

## Software availability

The whole data processing pipeline containing Python and R code is available in the form of Jupyter notebooks at https://github.com/canzarlab/heterochr_silencing

## Acknowledgments

The authors would like to thank Sigrun Jaklin for excellent technical assistance. This work was supported by St. Jude Children’s Research Hospital (M.H.), the American Lebanese Syrian Associated Charities (M.H.), ERC-smallRNAhet-309584 (M.H.), DFG HA 6941/6-1 (M.H.), NIH awards 1R01GM141694-01 (M.H.) and 1R01GM135599-01 (M.H.). P.M and P.K were supported by a Deutsche Forschungsgemeinschaft fellowship through the Graduate School of Quantitative Biosciences Munich.

## Author contributions

C.B and M.H. designed the experiments. C.B. and H.A. performed biochemical experiments. P.M.M, P.K. and C.B. performed bioinformatic analysis. M.H. and S.C. supervised the work. C.B., P.M.M, P.K., C.S. and M.H. wrote the paper.

## Methods

### Strain construction

All S. pombe strains used in this study are listed in **Supplemental Table S1**. The strains were constructed by electroporation (Biorad MicroPulser program ShS) with a PCR-based gene targeting product leading to deletion of specific genes. For genomic integration, a PCR with long overhang primers according to Bähler et al. (Bähler et al., 1998) was performed and the product transformed. Positive transformants were selected on YES plates containing 100 – 200 μg/ml antibiotics and were confirmed by PCR and sequencing. All *S. pombe* strains used in this study are listed in Supplemental Table S1. Strains were generated like described in (Brönner et al., 2017).

### total RNA isolation

Total RNA was isolated of a 2 ml yeast culture with OD 600 of 1.0 applying the hot phenol method. The pellet was resuspended in 500 μl lysis buffer (300 mM NaOAc pH 5.2, 10 mM EDTA, 1% SDS) and 500 μl phenol-chloroform-isoamylalcohol (25:24:1, Roth) and incubated at 65°C for 10 min with constant mixing. The organic and aqueous fractions were separated by centrifugation at 20 000 x g for 10 min. Nucleic acids in the aqueous fraction were precipitated with ethanol and then treated with DNAse I (Thermo Scientific) for 1 h at 37°C. DNAse was removed by a second phenol-chloroform-isoamylalcohol extraction and ethanol precipitation.

### poly(A) RNA sequencing

The poly(A) RNA library was obtained using the NEBNext Ultra II Directional RNA Library Prep Kit for Illumina (NEB) including the NEBNext Poly(A) mRNA Magnetic Isolation Module.

### Chromatin immunoprecipitation sequencing (ChIP-seq)

50 ml yeast cultures with an OD600 of 1.2 were cross-linked with 1% formaldehyde (Roth) for 15 min at room temperature. The reaction was quenched with 125 mM glycine for 5 min. The frozen pellet was resuspended in 500 µl lysis buffer (250 mM KCl, 1x Triton-X, 0.1% SDS, 0.1% Na-Desoxycholate, 50 mM HEPES pH 7.5, 2 mM EDTA, 2 mM EGTA, 5 mM MgCl2, 0.1% Nonidet P-40, 20% Glycerol) with 1 mM PMSF and Complete EDTA free Protease Inhibitor Cocktail (Roche). Lysis was performed with 0.25-0.5 mm glass beads (Roth) and the BioSpec FastPrep-24 bead beater (MP-Biomedicals), 8 cycles at 6.5 m/s for 30s and 3 min on ice. DNA was sheared by sonication (Bioruptor, Diagenode) 35 times for 30 s with a 30 s break. Cell debris was removed by centrifugation at 13 000 x g for 15 min. The crude lysate was normalized based on the RNA and Protein concentration (Nanodrop, Thermo Scientific) and incubated with 1.2 µg immobilized (Dynabeads Protein A, Thermo Scientific) antibody against Anti-RNA polymerase II CTD repeat YSPTSPS (phospho S2) antibody - ChIP Grade (ab5095, abcam) for at least 2 h at 4°C. The resin with immunoprecipitates was washed five times with each 1 ml of lysis buffer and eluted with 150 µl of elution buffer (50 mM Tris HCl pH 8.0, 10 mM EDTA, 1% SDS) at 65°C for 15 min. Cross-linking was reversed at 95°C for 15 min and subsequent RNase A (Thermo Scientific) digest for 30 min followed by Proteinase K (Roche) digest for at least 2 h at 37°C. DNA was recovered by phenol-chloroform-isoamylalcohol (25:24:1, Roth) extraction with subsequent ethanol precipitation. For sequencing, a ChIP-seq library was made using the NEBNext Ultra II DNA Library Prep Kit for Illumina kit (NEB).

### Pol II bound nascent RNA sequencing (RIP-seq)

RNA IP was performed like ChIP but without RNase A digest, using Anti-RNA polymerase II CTD repeat YSPTSPS (phospho S2) antibody - ChIP Grade (ab5095, abcam). After phenol-chloroform-isoamylalcohol extraction, DNA was digested with DNAse I (Thermo Scientific) for 2 h at 37°C. RNA was recovered with a second phenol-chloroform-isoamylalcohol purification and ethanol precipitation. Sequencing libraries were produced using the NEBNext Ultra II Directional RNA Library Prep Kit for Illumina (NEB). Libraries were sequenced on Illumina HiSeq platform.

### Analysis of sequencing data

Sequencing reads obtained in the poly(A) RNA sequencing (pA RNA), Pol II ChIP, nascent RNA sequencing (Pol II RIP) and total RNA sequencing experiments were mapped to the S. pombe reference genome (PomBase, release 2018) using splice-aware alignment tool STAR version 2.7.3a (Dobin et al., 2013). Alignment of RIP-seq and RNA-seq data was performed with STAR default parameters. Unspliced alignments of ChIP-seq data was enforced through parameters ‘--alignIntronMax 1’ and ‘--alignEndsType EndToEnd’. Reads mapping to ribosomal RNA have been removed from further analysis.

Genomic read counts were obtained using a custom script that extended basic htseq-count functionality with ‘--mode intersection-strict’ option. Additionally, for RNA assays pA RNA, Pol II RIP and total RNA, we used the ‘--stranded yes’ option to disambiguate which strand reads originated from. In short, reads mapping uniquely to protein-coding genes and reads that map less than 16 times were counted and normalized by gene length and sequencing depth (TPM, Transcripts Per Million). We chose 16 (*dg/dh* have in 12-13 copies) as multi-mapping threshold for heterochromatic genes to eliminate low-complexity reads without discarding reads originating from heterochromatic regions. In the read counting, we did not distinguish between individual gene copies but used one representative copy to collect all reads stemming from any one copy. Similarly, we obtained normalized read counts for intronic regions and defined the splice ratio for a given gene as the number of intronic reads divided by the corresponding genomic reads (Diaz et al., 2012). Average counts for protein coding and heterochromatic genes were computed from biological replicates which showed a high Spearman correlation score (> 80%) across all genes.

### INPUT subtraction in ChIP-seq data

For ChIP-seq data, separate scaling factors for IP-Input subtraction were computed for the 3 core centromeric regions such that the average background noise was close to zero. We selected these regions for normalization because they are the longest regions with very low transcription, therefore the signal present is defined as noise. In short, we computed coverages for ChIP-seq (x_i) and INPUT (y_i) for the selected regions and computed the INPUT normalization factor (lambda) using the following equation:

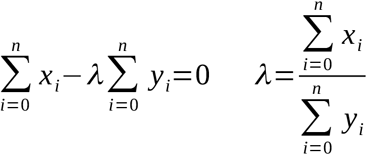

We used the median of the different values obtained for lambda as our final scaling factor. In Figure 1A we show that this transformation reduces notably the read counts associated to these regions in WT.

### Transcription Efficiency, RNA Stability, and Pol II Occupancy

For each gene, we measure transcription efficiency by the amount of newly synthesised RNA (Pol II Rip) relative to the level of Pol II Occupancy (Pol II ChIP):

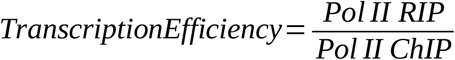

RNA levels being the result of RNA synthesis and degradation, we measure RNA stability as the ratio of steady state RNA level (pA RNA-seq) over RNA synthesis (Pol II RIP):

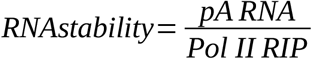

Change in PolII occupancy for the genes in each mutant was calculated as the log fold change of Pol-II occupancy as measured by Pol II Chip ChIP relative to wild type:

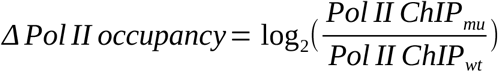

### Calculation of Contribution to silencing

First we calculated the enrichment of each pathway relative to wild type. We used the mean of each specific group (protein coding, *dg*/*dh*, *tlh*/*SPAC212.10*, heterochromatic genes) and then the mean of at least two duplicates. We calculated the enrichment of each pathway and each mutant over wild type. Quantification of each pathway’s contribution to heterochromatic silencing was calculated relative to *clr4Δ*. Additionally, wild type was set as 100 % silencing. We determined silencing according to the following equations :

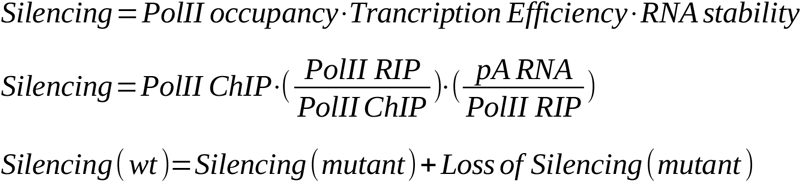

## Supplemental Information

**Supplemental Figure S1:**
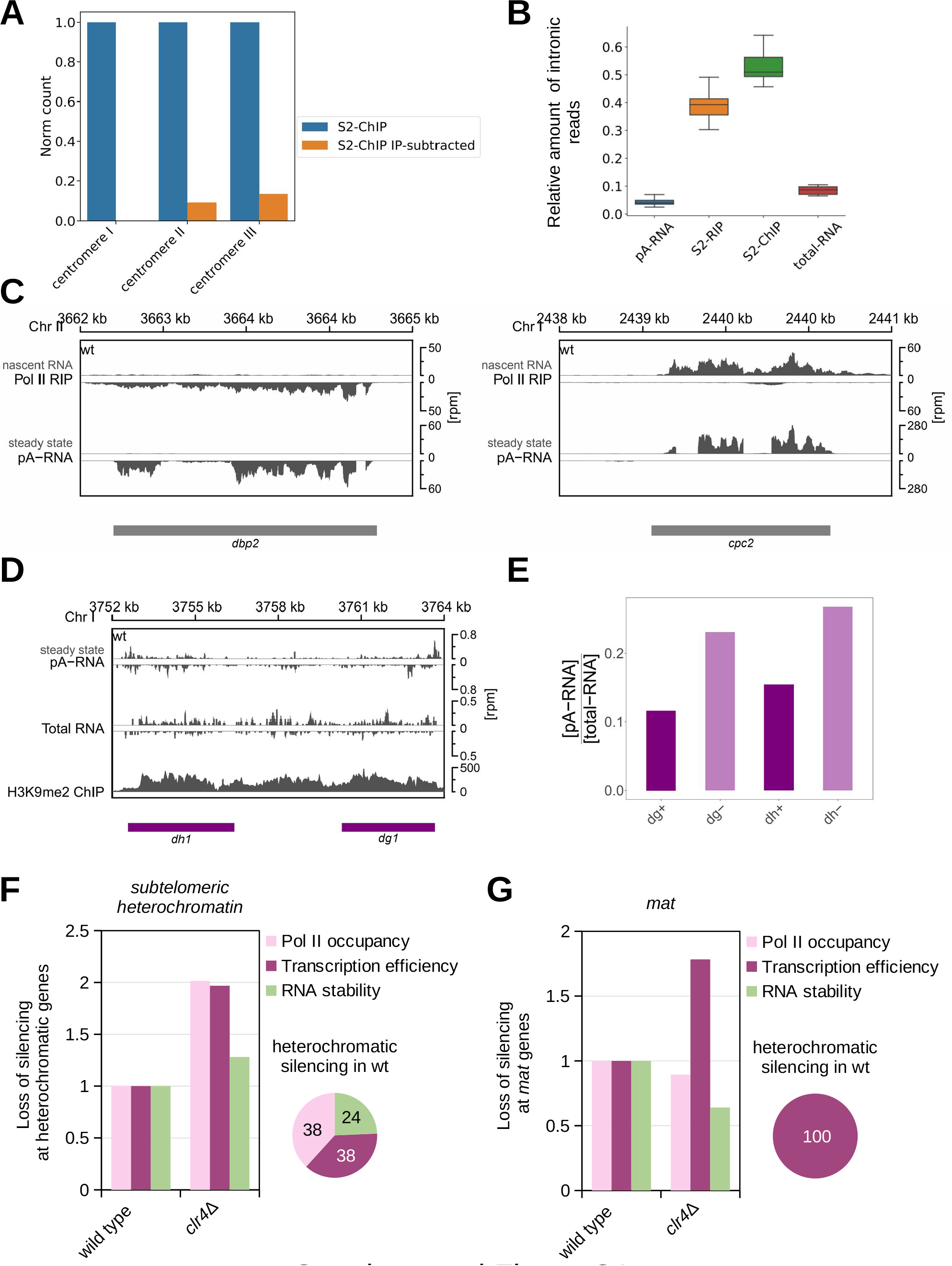
Silencing in wild-type cells. **(A)** Analysis of S2P-Pol II ChIPseq read counts from wild type cells at centromeric central core (Cenp-A chromatin) before and after normalized input subtraction. Note, after subtraction most reads from this region, defined as noise, are removed. **(B)** Quantification of intronic reads in pA RNA, total RNA, Pol II RIP and Pol II ChIP data. The data reveal high retention of intronic reads in Pol II RIP data. Bottom and top of the box correspond to lower and upper quartiles of the data, bar is the median and whiskers are median +/-1.5 times interquartile range. **(C)** Analysis of the next-generation sequencing data showing comparison between nascent RNA (RIP) and total RNA (pA) sequencing. Note presence of intronic reads in Pol II RIP data, indicating nascent RNAs. **(D)** Analysis of the next-generation sequencing data showing steady state RNA levels (total RNA-seq and pA RNA seq) and H3K9me2 levels (ChIP-seq) at pericentromeric regions in *S. pombe* wild-type cells. Locations of genes are indicated as boxes below the coverage according the color code: gray = protein coding; pink = *dg, dh*; green = *tlh, SPAC212.10*, blue = other heterochromatic genes. **(D)** Quantification of total and pA RNA reads over *dg* and *dh* transcripts showing that *dg+/dg-* and *dh+/dh-* transcripts are similarly polyadenylated. (**F, G**) Loss of silencing of the three pathways Pol II occupancy (S2P-Pol II ChIP), transcription efficiency (Pol II RIP / Pol II ChIP) and RNA stability (pA RNA / Pol II RIP) in *clr4Δ* cells at subtelomeric genes and *mat* locus. Pie chart demonstrates contribution of heterochromatic silencing in wild-type cells. Average of at least two independent samples is shown.

**Supplemental Figure S2:**
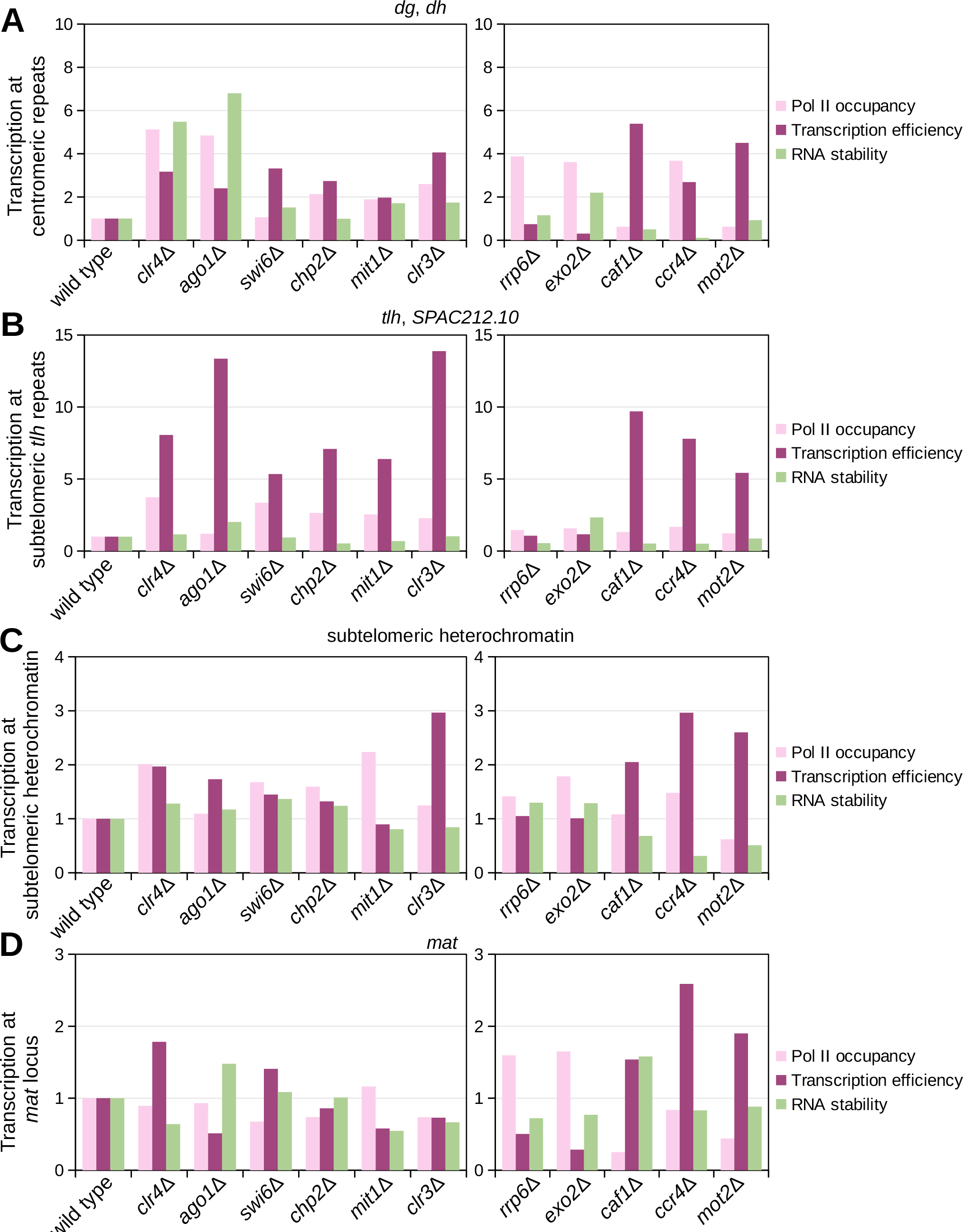
Silencing in mutant cells. (**A**) Loss of silencing at centromeric repeats in indicated mutant cells relative to wild type. Loss of silencing is shown for each of the three pathways RNA Pol II occupancy (Pol II ChIP), transcription efficiency (Pol II RIP / Pol II ChIP) and RNA stability (pA RNA / Pol II RIP). (**B**) Loss of silencing at subtelomeric *tlh* repeats in indicated mutant cells relative to wild type. Loss of silencing is shown for each of the three pathways RNA Pol II occupancy (Pol II ChIP), transcription efficiency (Pol II RIP / Pol II ChIP) and RNA stability (pA RNA / Pol II RIP). (**C**) Loss of silencing of the three pathways Pol II occupancy (Pol II ChIP), transcription efficiency (Pol II RIP / Pol II ChIP) and RNA stability (pA RNA / Pol II RIP) in several mutants at other subtelomeric genes. (**D**) Loss of silencing of the three pathways Pol II occupancy (Pol II ChIP), transcription efficiency (Pol II RIP / Pol II ChIP) and RNA stability (pA RNA / Pol II RIP) in several mutants at *mat* locus.

**Supplemental Figure S3:**
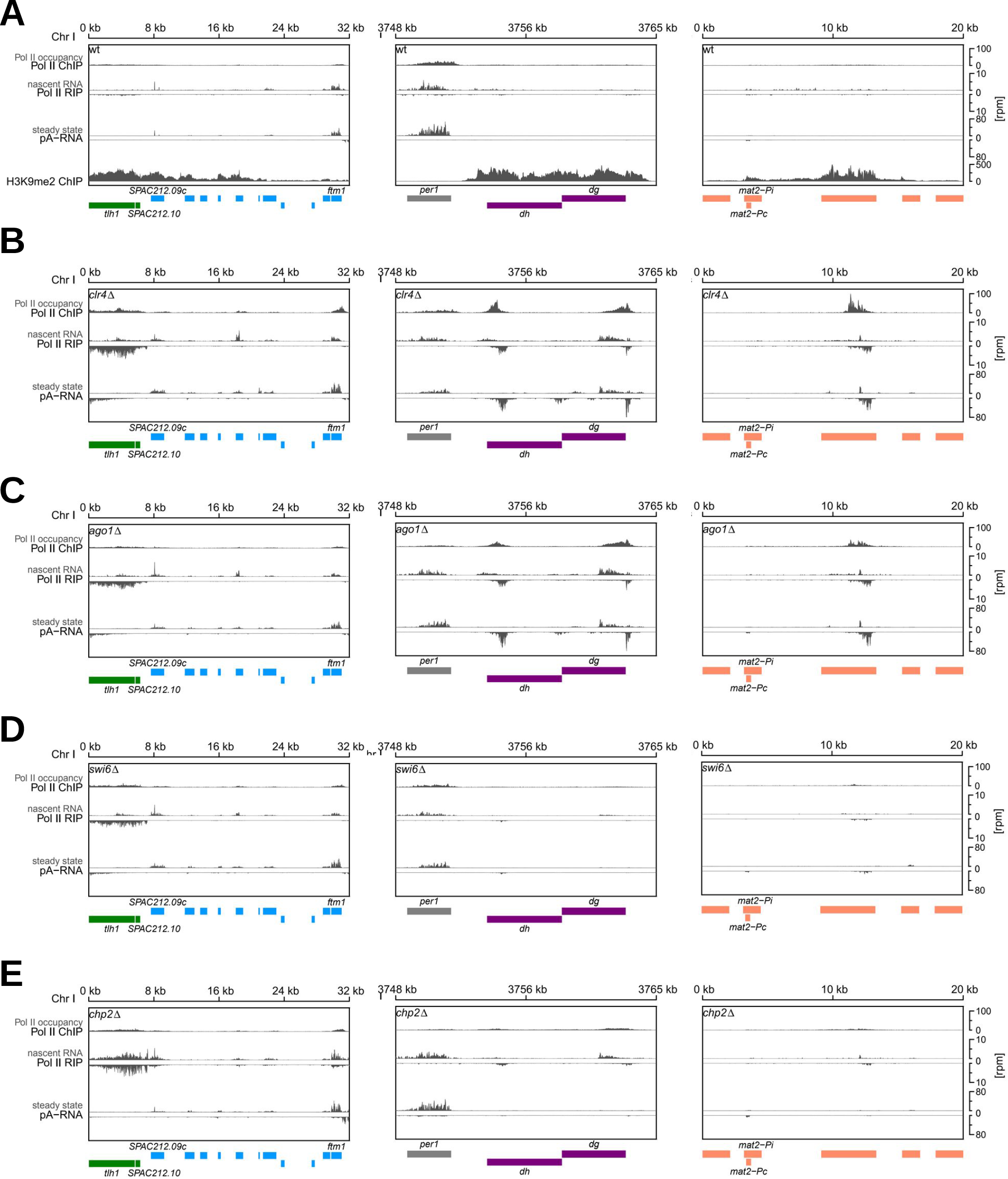
Next-Generation-Sequencing analysis of S2P-Pol II ChIP-seq (Pol II occupancy), S2P-Pol II RIP-seq (nascent RNA) and pA RNA-seq (steady state RNA) at subtelomeric and centromeric repeats, and at the *mat* locus. Locations of genes are indicated as boxes below the coverage according the color code: gray = protein coding; pink = *dg, dh*; green = *tlh, SPAC212.10*, blue = other heterochromatic genes. (**A**) wild type (**B**) *clr4Δ* (**C**) *ago1Δ* (**D**) *swi6Δ* (**E**) *chp2Δ*

**Supplemental Figure S4:**
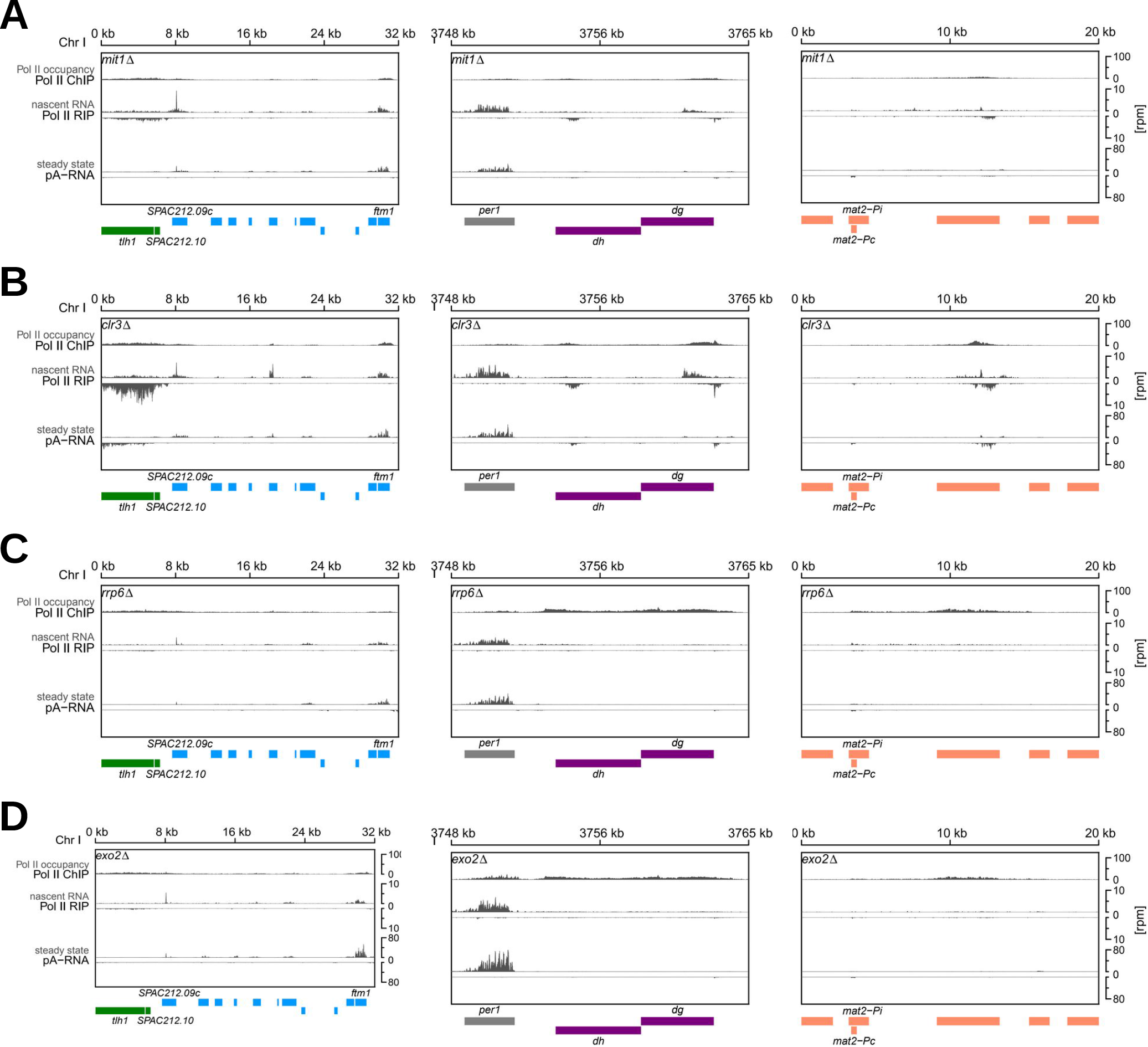
Next-Generation-Sequencing analysis of S2P-Pol II ChIP-seq (Pol II occupancy), S2P-Pol II RIP-seq (nascent RNA) and pA RNA-seq (steady state RNA) at subtelomeric and centromeric repeats, and at the *mat* locus. Locations of genes are indicated as boxes below the coverage according the color code: gray = protein coding; pink = *dg, dh*; green = *tlh, SPAC212.10*, blue = other heterochromatic genes. (**A**) *mit1Δ* (**B**) *clr3Δ* (**C**) *rrp6Δ* (**B**) *exo2Δ*

**Supplemental Figure S5:**
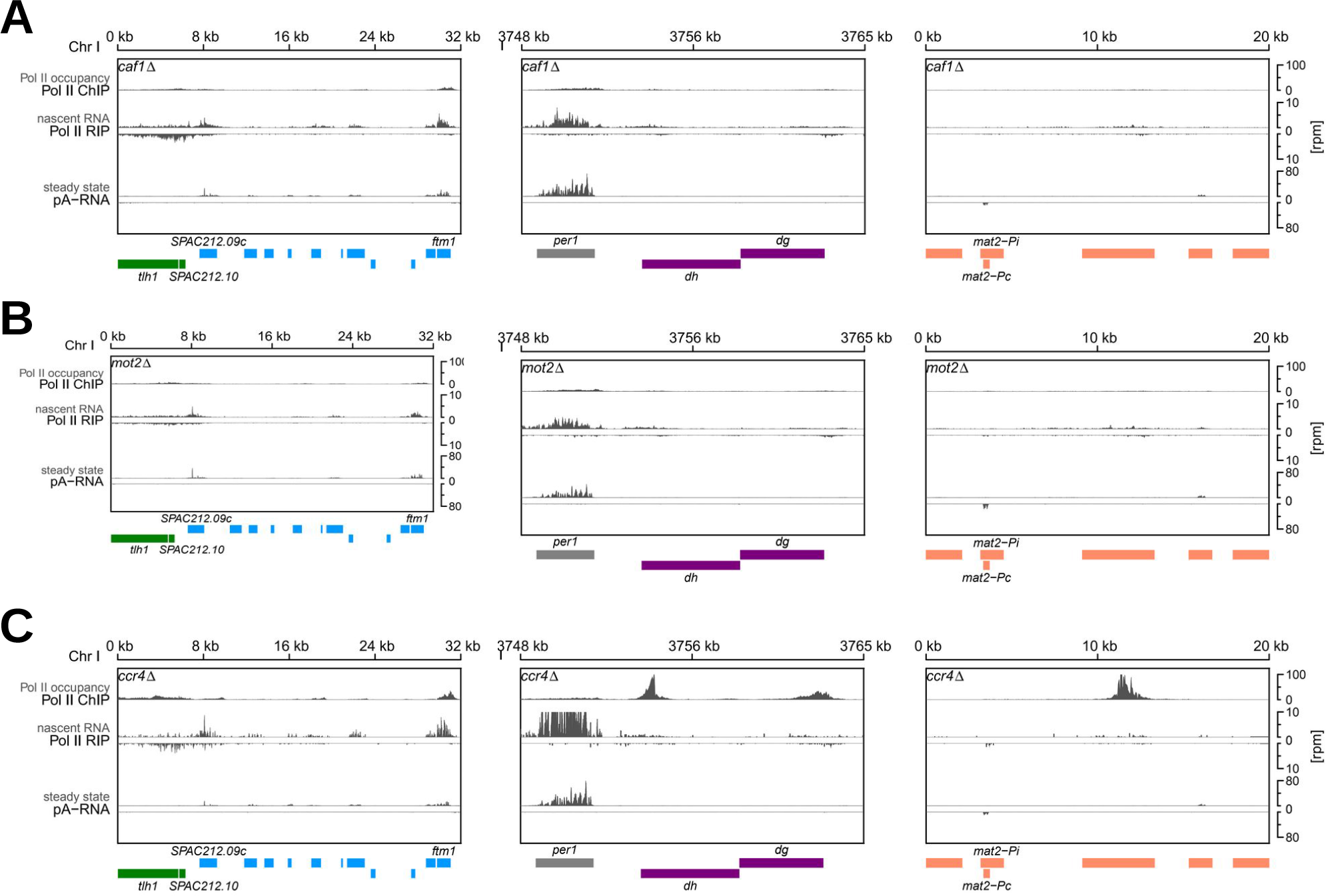
Next-Generation-Sequencing analysis of S2P-Pol II ChIP-seq (Pol II occupancy), S2P-Pol II RIP-seq (nascent RNA) and pA RNA-seq (steady state RNA) at subtelomeric and centromeric repeats, and at the *mat* locus. Locations of genes are indicated as boxes below the coverage according the color code: gray = protein coding; pink = *dg, dh*; green = *tlh, SPAC212.10*, blue = other heterochromatic genes. **(A)** *caf1Δ* **(C)** *mot2Δ* **(D)** *ccr4Δ*

**Supplemental Figure S6:**
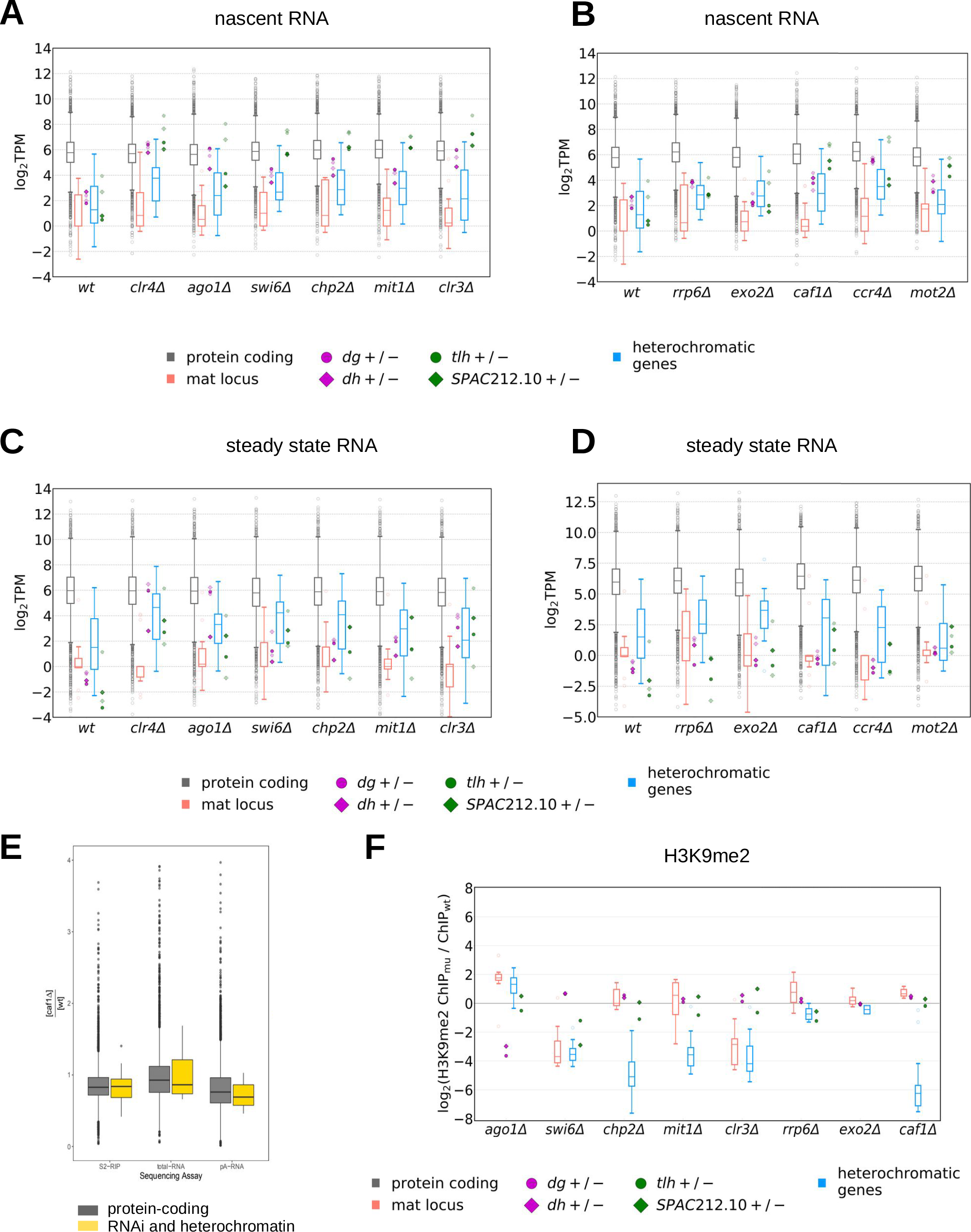
**(A, B)** Box plot showing S2P-Pol II RIP-seq data in wild-type and mutant cells. Nascent RNA analysis is shown for individual mutants affecting (A) heterochromatin formation or (B) RNA degradation. Data are plotted as defined for Figure 1C. (**C, D**) pA RNA-seq results (steady state RNA) shown as box plot for individual wild-type and mutants affecting (**C**) heterochromatin formation or (**D**) RNA degradation. Data are plotted as defined for Figure 1C. (**E**) Box plot showing ratio of RNA levels in S2P-Pol II RIP-seq, total RNA and pA RNA data. Protein coding genes are shown in grey and genes involved in RNAi and heterochromatin formation are shown in yellow. (**F**) Box plot showing H3K9me2 ChIP-seq data. H3K9me2 analysis is shown for individual mutants affecting heterochromatin formation or RNA degradation. Data are plotted as defined for Figure 1C. Average of at least two independent samples is shown.

**Supplemental Figure S7:**
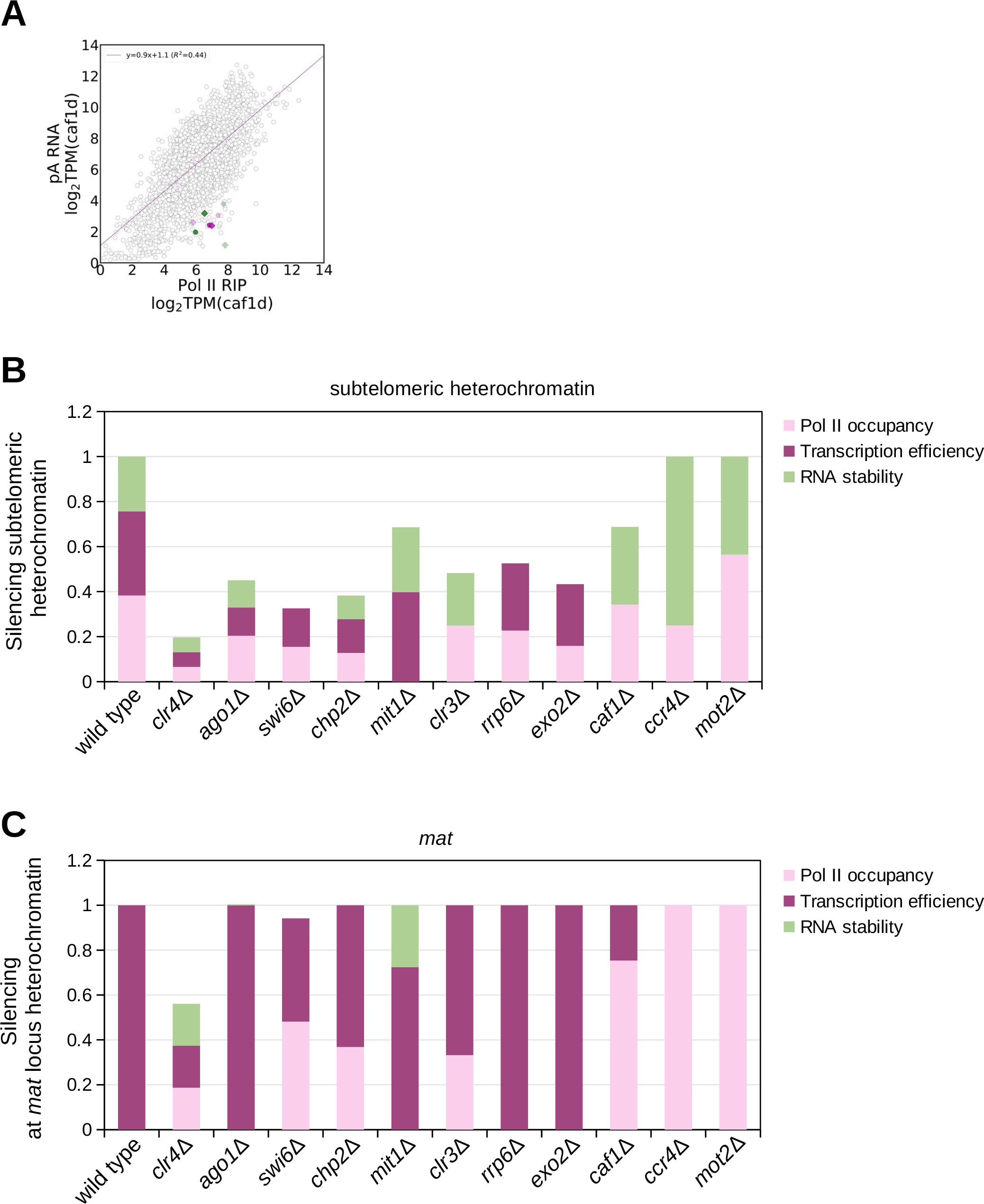
(**A**) RNA stability in *caf1Δ* cells. pA RNA-seq (steady state RNA) data plotted over S2P-Pol II RIP-seq data (nascent RNA). TPM, transcripts per million. Gray circles are individual protein-coding genes; regression line is also shown in gray. Also plotted are centromeric *dg* and *dh* (dark purple for + strand, bright purple for - strand) and *tlh* and *SPAC212.10* (dark green for + strand, bright green for - strand). Each data point is the average of at least two independent samples. (**B**) Bar chart displaying contribution of each pathway that is still active in the mutants to the silencing of other subtelomeric genes. (**C**) Bar chart displaying contribution of each pathway that is still active in the mutants to the silencing at the *mat* locus silencing.

**Supplemental Table S1:**
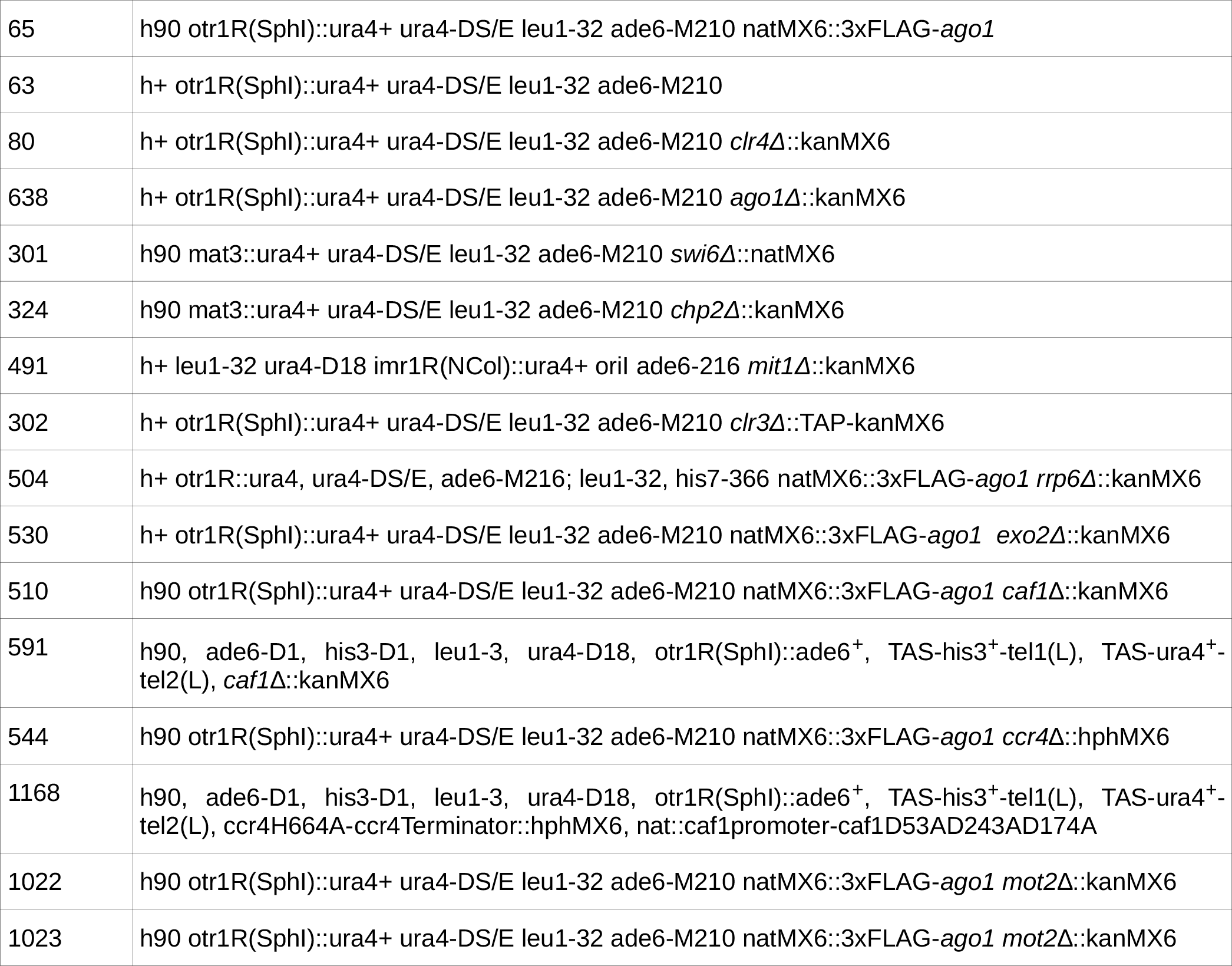
Strains used in this study.

**Supplemental Table S2:**
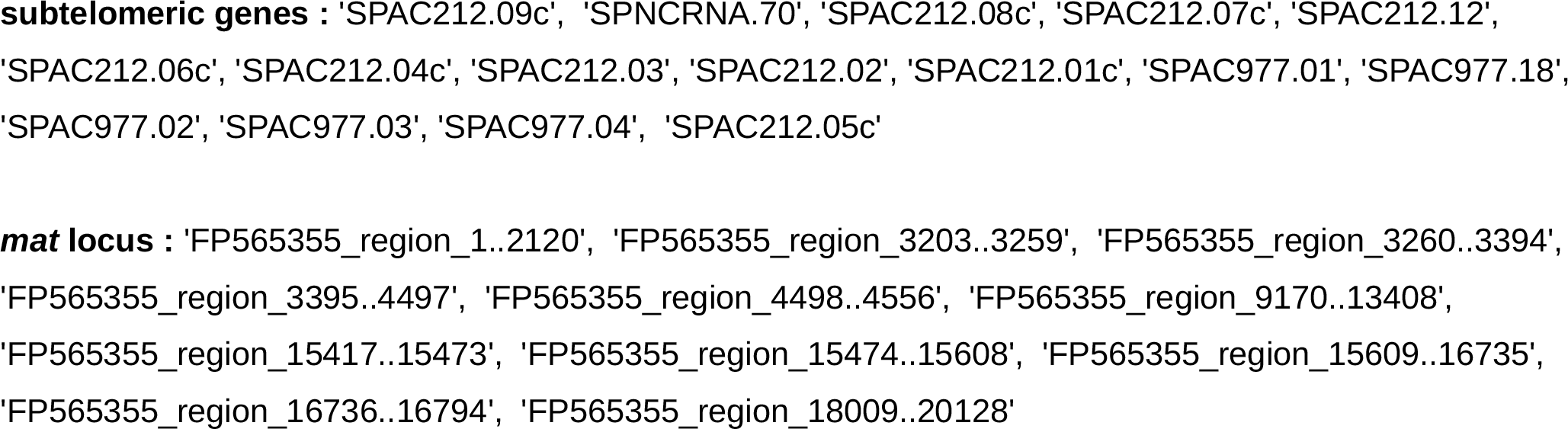
List of heterochromatic genes.

